# Rail-dbGaP: analyzing dbGaP-protected data in the cloud with Amazon Elastic MapReduce

**DOI:** 10.1101/035287

**Authors:** Abhinav Nellore, Christopher Wilks, Kasper D Hansen, Jeffrey T Leek, Ben Langmead

## Abstract

**Motivation**: Public archives contain thousands of trillions of bases of valuable sequencing data. More than 40% of the Sequence Read Archive is human data protected by provisions such as dbGaP To analyze dbGaP-protected data, researchers must typically work with IT administrators and signing officials to ensure all levels of security are implemented at their institution. This is a major obstacle, impeding reproducibility and reducing the utility of archived data.

**Results**: We present a protocol and software tool for analyzing protected data in a commercial cloud. The protocol, Rail-dbGaP, is applicable to any tool running on Amazon Web Services Elastic MapReduce. The tool, Rail-RNA v0.2, is a spliced aligner for RNA-seq data, which we demonstrate by running on 9,662 samples from the dbGaP-protected GTEx consortium dataset. The Rail-dbGaP protocol makes explicit for the first time the steps an investigator must take to develop Elastic MapReduce pipelines that analyze dbGaP-protected data in a manner compliant with NIH guidelines. Rail-RNA automates implementation of the protocol, making it easy for typical biomedical investigators to study protected RNA-seq data, regardless of their local IT resources or expertise.

**Availability**: Rail-RNA is available from http://rail.bio. Technical details on the Rail-dbGaP protocol as well as an implementation walkthrough are available at https://github.com/nellore/rail-dbgap. Detailed instructions on running Rail-RNA on dbGaP-protected data using Amazon Web Services are available at http://docs.rail.bio/dbgap/.

## 1 INTRODUCTION

The Database of Genotypes and Phenotypes (dbGaP) (Mailman *et al*., 2007) hosts controlled-access raw and processed human genomic data and associated phenotypic data. While datasets are stripped of metadata linking them with specific individuals, the remaining data is still sensitive in part because it could be combined with external information to identify individuals. The NIH requires adherence to security guidelines for the proper handling of dbGaP-protected data, from acquisition through destruction (see http://www.ncbi.nlm.nih.gov/projects/gap/pdf/dbgap_;2b_security_procedures.pdf). These include physical security of computing infrastructure, restricting inbound internet access, multi-factor authentication (MFA) and password policies, encryption of data, enforcing the principle of least privilege, and logging data access. For many investigators, the guidelines pose a practical challenge: institutional computer clusters may not be compliant, requiring IT policy revisions or intervention by IT administrators.

The recent NIH announcement (https://grants.nih.gov/grants/ guide/notice-files/NOT-OD-15-086.html) allowing investigators to request permission to transfer to and analyze dbGaP data in compliant clouds provides a convenient alternative: investigators can use protocols and software tailored for secure analysis of dbGaP data in a commercial cloud, bypassing local infrastructure issues.

Commercial cloud providers allow users to rent computational and storage resources residing in large data centers. Reputable providers like Amazon, Microsoft, and Google ensure physical security of data centers. Resources come in standardized units of hardware and software; the hardware is rented in standard (usually virtualized) units called instances and software comes pre-installed on standard disk images selected at launch time.

Amazon Web Services (AWS) is a popular choice for genomics data analysis. Key datasets such as 1000 Genomes, TCGA, and ICGC are hosted in AWS storage, allowing researchers to use cloud-enabled tools without copying data across the internet (see https://aws.amazon.com/1000genomes/ and https://aws.amazon.com/public-data-sets/tcga/). Input data, intermediate results, and final results can all be kept in cloud storage before final results are downloaded or browsed. AWS also provides guidance on secure analysis of protected genomic data (see https://d0.awsstatic.com/whitepapers/compliance/AWS_dBGaP_Genomics_on_AWS_Best_Practices.pdf).

We describe two important first steps toward enabling typical investigators to analyze dbGaP-protected data on AWS: (1) Rail-dbGaP, a protocol for analyzing dbGaP data in the cloud using Elastic MapReduce (EMR), an AWS service; and (2) the v0.2 line of our Rail-RNA tool (Nellore *et al.*, 2015), which implements the protocol to securely download and align many dbGaP-protected RNA sequencing (RNA-seq) samples at once on EMR. We demonstrate the tool by running it on 9,662 RNA-seq samples from the GTEx project.

**Fig. 1.**
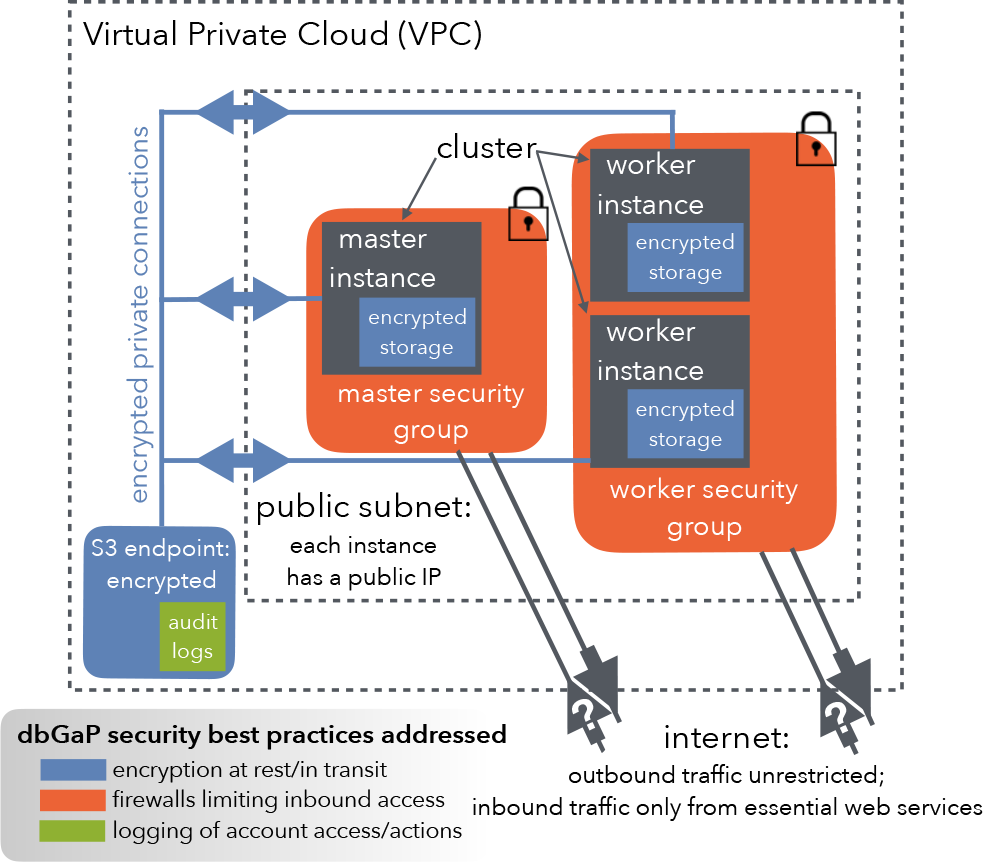
The Rail-dbGaP security architecture. Security features include a Virtual Private Cloud with a private connection between the computer cluster and cloud storage, audit logs recorded on cloud storage, encryption of sensitive data at rest and in transit, and restricted inbound access to the cluster via security groups.

## 2 PROTOCOL FEATURES

An EMR Hadoop (http://hadoop.apache.org) cluster consists of a master instance and several worker instances. The Rail-dbGaP security architecture (Fig. 1) secures the cluster to satisfy dbGaP guidelines. These guidelines are described below in the context of AWS and are formulated more precisely in the supplementary document entitled, “The Rail-dbGaP protocol,” also available at https://github.com/nellore/rail-dbgap/blob/master/README.md. The supplementary document further walks the reader through the development of an EMR pipeline that uses the Rail-dbGaP protocol to analyze three dbGaP-protected test samples.

- **Cluster is within a subnet of a Virtual Private Cloud (VPC)**. A VPC is a logically isolated unit of the cloud providing a private network and firewall. The connection with the cloud storage service (Amazon Simple Storage Service, or S3) is via a “VPC endpoint,” which ensures that data transferred never leaves the data center.
- **Inbound traffic is restricted via security groups**. A security group is essentially a stateful firewall. A master security group for the master instance and a worker security group for worker instances prevent initiation of any connection to the cluster except by essential web services.
- **Data are encrypted at rest**. During cluster setup, before any sensitive data has reached the cluster, each instance runs a preliminary script (“bootstrap action”) that uses Linux Unified Key Setup (LUKS) (https://guardianproject.info/code/luks/) to create an encrypted partition with a keyfile. The key is randomly generated on each instance and never exposed to the user. Temporary files, the Hadoop distributed file system, and buffered output to the cloud storage service are all configured to reside on the encrypted partition via symbolic links. Files written to cloud storage are also encrypted. On S3, this is enforced by creating a bucket policy (i.e., rules governing user access to the bucket) barring uploads that do not use server-side encryption.
- **Data are encrypted in transit**. Worker instances download dbGaP data using SRA Tools (http://ncbi.github.io/sra-tools/), ensuring encryption of data transferred from dbGaP to the cluster. Secure Sockets Layer (SSL) is enabled for transfers between cloud storage and the cluster as well as between cloud storage service and compliant local storage to which an investigator saves results.
- **Identities are managed to enforce the principle of least privilege**. The principle of least privilege prescribes users have only the privileges required to perform necessary tasks. In the Rail-dbGaP protocol, an administrator uses multi-factor authentication, grants the user only necessary privileges, and constrains the user to set up a password satisfying NIH security best practices at http://www.ncbi.nlm.nih.gov/projects/gap/pdf/dbgap2bsecurityprocedures.pdf.
- **Audit logs are enabled**. These record logins and actions taken by the user and on the user’s behalf, including API calls made by processes running on the cluster. On AWS, audit logs take the form of CloudTrail logs stored in encrypted S3 buckets.

## 3 APPLICATION

The Rail-dbGaP protocol is implemented in the v0.2 line of Rail-RNA. Detailed instructions on implementation are available at https://github.com/nellore/rail-dbgap, and Rail-RNA v0.2.x may be downloaded at http://rail.bio. We used the Rail-dbGaP protocol to align 9,662 paired-end RNA-seq samples obtained by the GTEx consortium (Lonsdale et al., 2013) in 30 batches across various human tissues over a period of 5 days, 5 hours, and 28 minutes for US$0.32 per sample. Computational details, mapped read proportions, and a cost calculation are described in the Supplementary Material. Scripts for reproducing our results are available at https://github.com/nellore/runs/tree/master/gtex.

## ACKNOWLEDGMENTS

We thank the GTEx project for making data available. We thank Christopher Goodson, Lenworth Henry, and Angel Pizarro of Amazon Web Services and Ronald Dowden and Brian Willey of IT@JH for assistance in designing and validating the protocol. Funding: AN, JTL, and BL were supported by NIH/NIGMS grant 1R01GM105705 to JTL. AN was supported by a seed grant from the Institute for Data Intensive Engineering and Science (IDIES) at Johns Hopkins University to BL and JTL. BL was supported by a Sloan Research Fellowship to BL. Amazon Web Services experiments were supported by AWS in Education Research grants.

## SUPPLEMENTARY MATERIAL

### 3.1 Computation details

We analyzed 9,662 RNA-seq samples from the V6 release by the GTEx consortium. These were divided randomly into 30 batches of approximately the same size: two batches had 323 samples, and the others had 322 samples. Rail-RNA splits analysis of each batch up into a preprocess job flow, which securely downloads raw reads from dbGaP and uploads preprocessed versions to S3; and an alignment job flow, which aligns preprocessed reads and writes results back to S3. Preprocess job flows were run on 21 m3.xlarge Amazon EC2 instances, each with four 2.4 GHz Intel Xeon E5-2670 v2 (Ivy Bridge) processing cores and 15 GB of RAM. Alignment job flows were run on 81 c3.8xlarge Amazon EC2 instances, each with 32 Intel Xeon E5-2680 v2 (Ivy Bridge) processing cores and 60 GB of RAM. One instance of every EMR cluster was a master and the rest were workers, so up to 80 worker processing cores were active for each preprocess job flow and up to 2560 worker processing cores were active for each alignment job flow. All EC2 instances were obtained from the EC2 spot marketplace, which allowed us to reduce our cost to the fluctuating market price, typically a fraction of the standard (on-demand) price. Job flows were submitted manually, with preprocess job flows staggered to minimize issues with simultaneous downloads from the dbGaP server. The number of rented worker cores across the 5 days, 5 hours, and 28 minutes during which job flows were run is summarized in Fig. 2. We typically could not download more than 240 samples at once across all our preprocess job flows without sustaining failures; jumps above 240 active cores in the activity for preprocess job flows depicted in Fig. 2 point to inactive cores on EMR clusters waiting for hanging tasks to complete. Rail-RNA users can expect to observe similar behavior. Mapped read proportions across GTEx are depicted in Fig. 3.

**Fig. 2.**
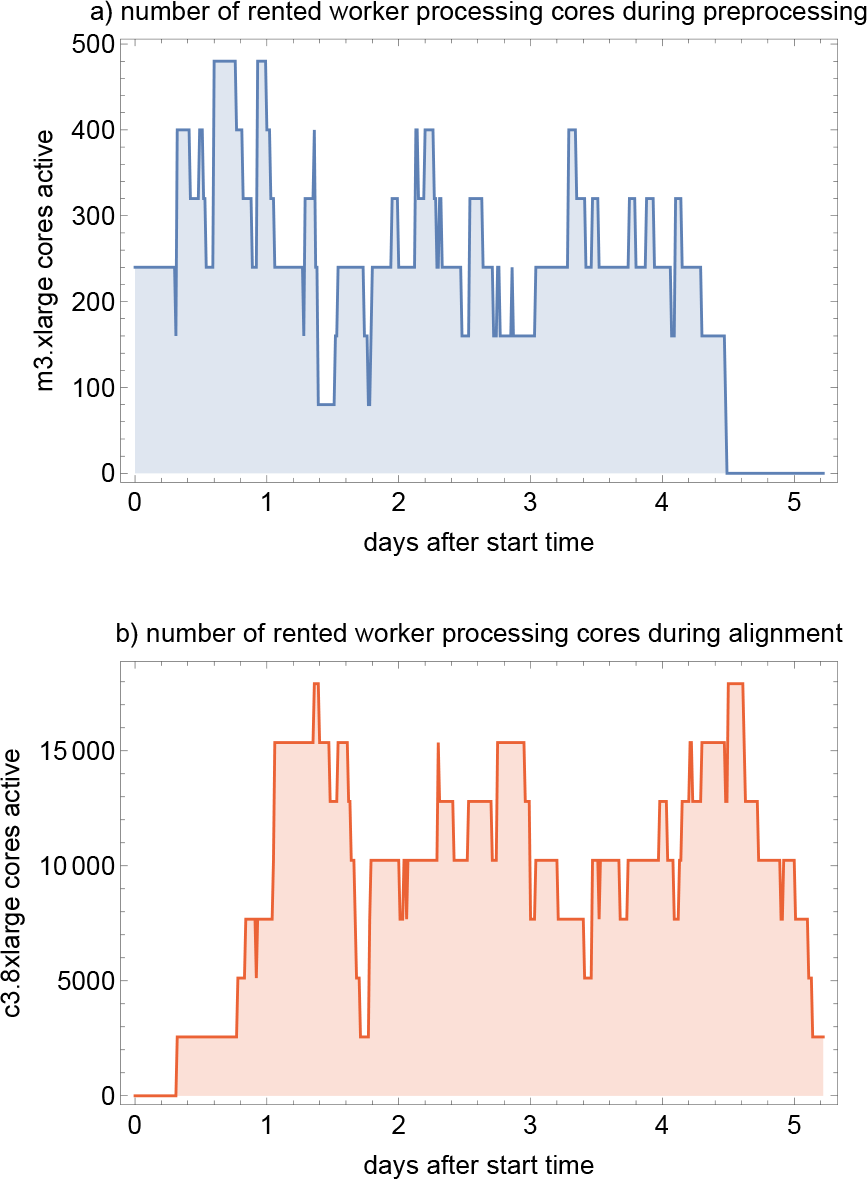
The number of rented worker cores across clusters running a) preprocess and b) alignment job flows for the duration of all job flows.

**Fig. 3.**
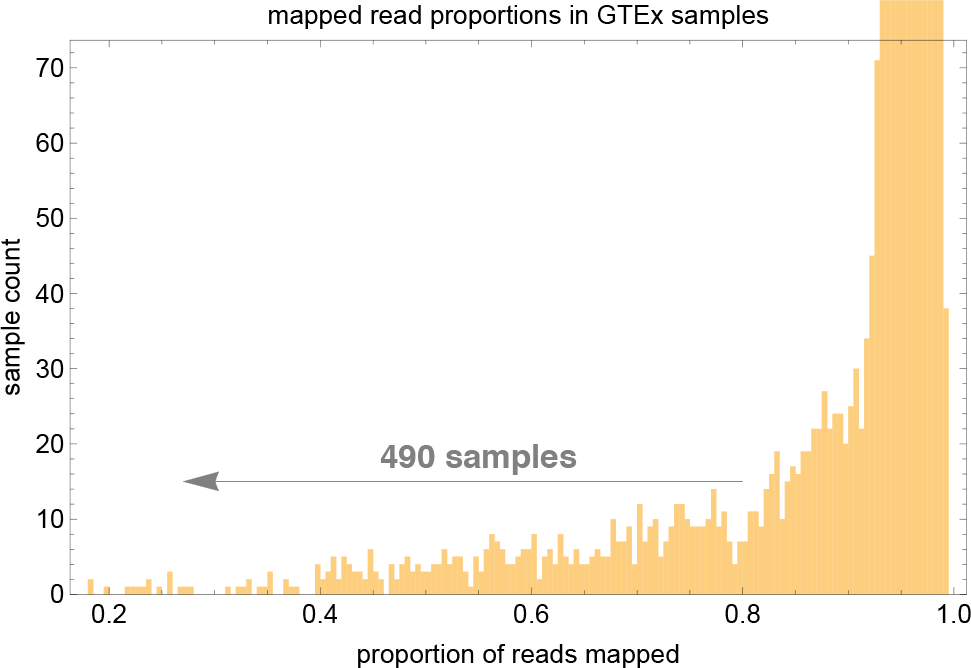
Mapped read proportions across 9,662 GTEx RNA-seq samples from Rail-RNA alignment job flows. 490 samples each had fewer than 80% mapped reads.

### 3.2 Cost calculations

We used the AWS Cost Explorer to sum costs across the five days over which the computation ran: November 30, 2015 through December 4, 2015. The total cost of our analysis was US$28,368.15, which gives an average cost of US$0.32 per 10 million reads aligned. Raw cost data downloaded from the AWS Cost Explorer are available at https://github.com/nellore/runs/blob/master/gtex/costs.csv. Costs broken down by AWS service are depicted in Fig. 4.

**Fig. 4.**
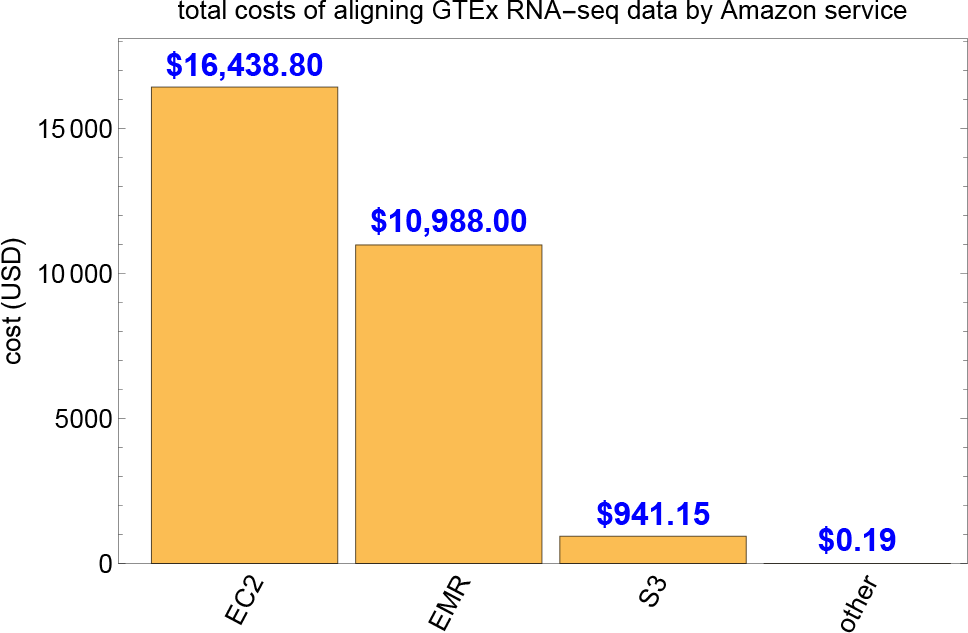
Total costs of GTEx analysis jobs divided up by Amazon service. The trivial contributions of the “other” costs are from Simple Queue Service (SQS) and SimpleDB.

